# Beyond mean trabecular separation: decomposing metaphyseal marrow space into local trabecular spacing and marrow cavity expansion

**DOI:** 10.64898/2026.06.17.729075

**Authors:** Carmen Huesa, John C. Lockhart, Carl S. Goodyear, Jonathan A. Williams

**Affiliations:** School of Infection and Immunity, College of Medical, Veterinary and Life Sciences, University of Glasgow, G12 8QQ, UK; Institute of Biomedical and Environmental Health Research, University of the West of Scotland; Department of Biomedical Engineering, Wolfson Building, University of Strathclyde, Glasgow, G4 0NW, UK

**Keywords:** trabecular bone, micro-computed tomography (µCT), bone morphometry, trabecular bone separation, osteoporosis, preclinical bone models

## Abstract

Micro-computed tomography (µCT) is widely used to assess trabecular bone microarchitecture, with trabecular separation (Tb.Sp) among the core parameters recommended for reporting. Tb.Sp is typically expressed as a single volume-weighted mean derived from maximal sphere fitting, although the underlying distribution of local separation values is rarely examined.

Here, we show that Tb.Sp distributions in metaphyseal trabecular bone are frequently non-Gaussian and bimodal or multimodal. Using µCT datasets from three established models of osteoporosis, spinal cord injury (SCI), ovariectomy (OVX), and ageing, we demonstrate that this behaviour is most evident in metaphyseal trabecular bone and is less apparent in epiphyseal trabecular bone or trabecular thickness distributions.

We further show that multimodal metaphyseal Tb.Sp distributions correspond to two spatially distinct contributions within the marrow space: lower-diameter local separation within the residual trabecular network, and higher-diameter regions associated with larger contiguous marrow cavities. Based on this observation, we introduce a simple extension to standard morphometric analysis in which Tb.Sp is decomposed into local trabecular separation (Tb.Sp_L_) and marrow cavity separation (Tb.Sp_M_).

Tb.Sp decomposition revealed model-specific patterns of trabecular deterioration. SCI was characterised predominantly by increased Tb.Sp_M_, consistent with expansion of larger marrow cavities, whereas OVX showed a more subtle or distributed alteration. Ageing showed changes in both Tb.Sp_L_ and Tb.Sp_M_, with the higher-diameter component becoming most prominent in older animals. Together, these findings demonstrate that mean Tb.Sp can mask structurally distinct forms of metaphyseal marrow-space organisation and support reporting distributional descriptors, and where appropriate Tb.Sp_L_ and Tb.Sp_M_, alongside conventional Tb.Sp.

## 1. Introduction

Structurally, osteoporosis is characterised by loss of bone mass and deterioration of bone microarchitecture, leading to reduced mechanical competence and increased fracture risk (LeBoff et al., 2022). Although bone mineral density is widely used to assess skeletal health, it does not fully capture the architectural changes that contribute to fracture resistance (Seeman & Delmas, 2006). Trabecular bone is particularly susceptible to remodelling imbalances because of its high surface-to-volume ratio (Koh et al., 2024), and degradation of trabecular-rich regions such as the spine, hip, and metaphyses of long bones is a key feature of osteoporotic bone loss (LeBoff et al., 2022). Accurate interpretation of these changes, therefore, requires morphometric approaches that capture not only the amount of trabecular bone present, but also how the trabecular architecture and marrow space are spatially organised.

Micro-computed tomography (µCT) is the gold-standard technique for assessing trabecular bone microarchitecture in preclinical models. Standardised guidelines for µCT-based morphometry recommend reporting a core set of trabecular parameters, including bone volume fraction (BV/TV), trabecular thickness (Tb.Th), trabecular number (Tb.N), and trabecular separation (Tb.Sp) (Bouxsein et al., 2010). These parameters provide quantitative descriptors of trabecular bone structure and are routinely used to characterise skeletal phenotypes across a wide range of experimental conditions.

In rodent models, trabecular bone morphometry is most commonly performed in the metaphyseal regions of long bones, particularly the distal femur and proximal tibia (Salmon et al., 2023). These sites are widely used because they contain relatively large volumes of trabecular bone in small animals, are highly responsive to experimental perturbations such as ovariectomy, ageing, and mechanical unloading, and have a long history of use in both histological and µCT-based bone morphometry (Salmon et al., 2023). They also offer practical advantages for image analysis: within the secondary spongiosa, trabecular bone can usually be separated from the surrounding cortical bone shell using reproducible semi-automated or automated methods (Buie et al., 2007). This is more challenging in geometrically complex regions such as the epiphysis, where the trabecular compartment is less straightforward to isolate (Herbst et al., 2021).

The metaphysis, however, should not be regarded as a uniform structural compartment. It extends from the growth plate toward the diaphysis and contains gradients in trabecular density, organisation, and maturational state, including the transition from primary to secondary spongiosa (Erben, 1996; Salmon et al., 2023). Previous work has shown that resolving metaphyseal trabecular bone profiles as a function of distance from the growth plate can reveal information that is obscured by whole volume of interest (VOI) averages (Gabet et al., 2008; Williams et al., 2020; Salmon et al., 2023). This is particularly important in metaphyseal bone, where the interpretation of a morphometric parameter depends not only on the calculation performed, but also on the VOI over which it is measured. A standard metaphyseal VOI may include both dense residual trabecular networks and larger marrow-space regions. Whole-VOI averages can therefore combine structurally distinct features into a single value, making metaphyseal morphometry informative but potentially difficult to interpret using summary parameters alone.

Trabecular separation (Tb.Sp) is particularly relevant in this context because it describes the marrow-space phase rather than the bone phase. Tb.Sp estimates the local thickness of the marrow space between trabeculae and is generally calculated using a model-independent 3D local-thickness approach (Hildebrand & Rüegsegger, 1997). In this formulation, the local thickness at a point is defined as the diameter of the largest sphere that contains that point and remains entirely bounded within the analysed phase; the point is not necessarily located at the centre of the sphere (Hildebrand & Rüegsegger, 1997). Conventional reporting of Tb.Sp collapses this local-thickness field into a single volume-weighted mean value. However, the same calculation can also be represented as a voxel-wise map (Figure 1A), a histogram of local separation values (Figure 1B), or distributional summary statistics such as the standard deviation, SD(Tb.Sp), or maximum local separation value, max(Tb.Sp) (Salmon, 2020; Domander et al., 2021).

**Figure 1.**
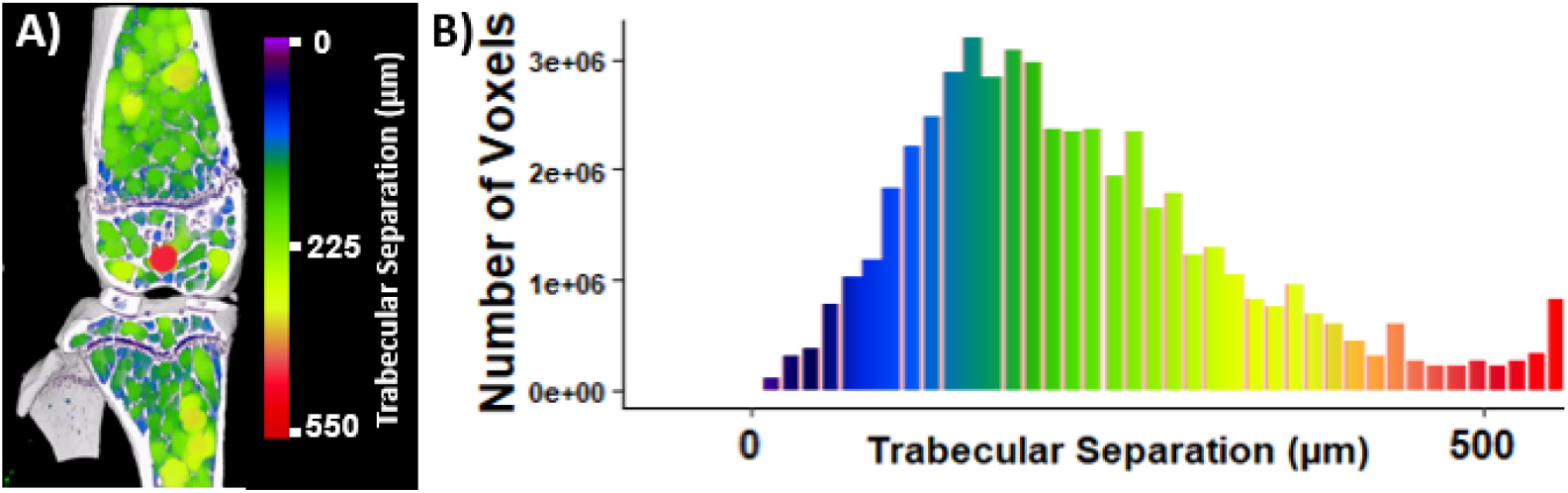
Example voxel-wise trabecular separation (Tb.Sp) map and histogram. **A)** Representative voxel-wise Tb.Sp map from a mouse long-bone µCT dataset, shown across the knee region, including proximal tibia and distal femur. Tb.Sp was calculated as the local thickness of the marrow-space phase, in which each marrow space voxel is assigned a local separation value corresponding to the diameter of the largest sphere that contains that voxel and remains bounded within the marrow space. Colours encode local Tb.Sp, according to the adjacent rainbow scale, with larger values shown toward the red end of the scale. The binarised bone is shown in white. **B)** Histogram of voxel-wise Tb.Sp values from the map shown in A. Bar height represents the number of voxels within each Tb.Sp range, and bar colour corresponds to the Tb.Sp colour scale in A.

Although the mean Tb.Sp is widely reported, it remains unclear how much structural information is lost when the voxel-wise marrow-space thickness field is reduced to a single summary value. Existing approaches, including distance-from-growth-plate profiling, additional distributional descriptors, and architectural or topological measures such as connectivity (Odgaard & Gundersen, 1993), anisotropy (Harrigan & Mann, 1984), fractal dimension (Fazzalari & Parkinson, 1996), ellipsoid factor (Doube, 2015) and inter-trabecular angle (Reznikov et al., 2016), provide valuable methods to quantify heterogeneity in trabecular bone architecture. However, these approaches generally describe spatial variation, network organisation, or global structural complexity, rather than directly analysing whether the voxel-wise marrow-space thickness distribution itself contains distinct modal components. Although Tb.Sp distributions have been reported previously (Liu et al., 2015; Kerckhofs et al., 2016; Porrelli et al., 2022), whether such components can be separated within a single metaphyseal VOI remains unclear.

In this study, we tested whether Tb.Sp distributions in metaphyseal trabecular bone contain distinct lower- and higher-diameter components that can be identified from the voxel-wise Tb.Sp map and corresponding histogram. We introduce a simple extension to standard morphometric analysis in which Tb.Sp is decomposed into local trabecular separation, Tb.Sp_L_, and marrow cavity separation, Tb.Sp_M_. To evaluate this approach, we analysed µCT datasets from three established models of osteoporosis representing distinct mechanisms of bone loss: spinal cord injury-induced mechanical unloading, ovariectomy-induced hormone deficiency, and ageing. Specifically, we aimed to: (i) characterise the distributional behaviour of metaphyseal Tb.Sp across independent osteoporosis datasets; (ii) determine whether bimodal Tb.Sp distributions correspond to spatially distinct regions within the metaphyseal marrow space; (iii) assess whether decomposition into Tb.Sp_L_ and Tb.Sp_M_ provides information beyond the conventional mean Tb.Sp; and (iv) examine whether the relative behaviour of Tb.Sp_L_ and Tb.Sp_M_ varies between osteoporosis models with distinct biological origins.

## 2. Methods

### 2.1 µCT datasets, VOI definition, and segmentation

Micro-computed tomography (µCT) datasets from three independent small-animal studies were re-analysed: an unpublished ovariectomy-induced osteoporosis (OVX) study in mice, a published ageing (AGE) study in mice (Huesa et al., 2026), and published spinal cord injury-induced osteoporosis (SCI) studies in rats (Williams et al., 2020, 2022). Together, these datasets represent distinct biological mechanisms of bone loss: hormone deficiency, ageing, and mechanical unloading, respectively. Metaphyseal trabecular bone was analysed in all datasets, comprising proximal tibial metaphyseal trabecular bone for the OVX and AGE datasets and distal femoral metaphyseal trabecular bone for the SCI dataset.

Metaphyseal VOIs were defined relative to the growth plate in each dataset, using an offset selected to exclude primary spongiosal structures. VOIs were standardised within each study and were not adjusted to the reduced trabecular extent in the osteoporotic groups. This retained reductions in trabecular extent and associated expansion of the metaphyseal marrow space within the analysis.

Datasets were binarised using study-specific global thresholds to separate mineralised tissue from non-mineralised space. Within each metaphyseal VOI, trabecular bone was then isolated from the surrounding cortical bone using standard automated methods. The resulting binary trabecular VOIs were used for all subsequent morphometric analyses, with the marrow-space phase for Tb.Sp analysis defined as the inverse of the binary trabecular bone phase within each VOI.

As the datasets were derived from independent studies, acquisition parameters, VOI dimensions, and segmentation procedures varied between datasets. Specimen characteristics, scan settings, reconstruction settings, VOI definitions, and dataset-specific processing details are summarised in Supplementary Table S1.

### 2.2 Standard morphometric analysis

Standard 3D trabecular morphometric parameters, including BV/TV, Tb.Th, Tb.N, and Tb.Sp, were quantified from the segmented datasets in CTAnalyzer software (CTAn) (version 1.23.02, Bruker, Belgium) following established µCT bone morphometry guidelines (Bouxsein et al., 2010).

For Tb.Sp analysis, CTAn generated a Tb.Sp histogram (Figure 1B), from which summary morphometric values such as the volume-weighted mean Tb.Sp and SD(Tb.Sp) were derived, together with a voxel-wise Tb.Sp map (Figure 1A). In CTAn, Tb.Sp histograms are reported using fixed bin intervals corresponding to successive local-separation ranges of 1 – <3, 3 – <5, 5 – <7, and so forth, expressed in calibrated units based on the dataset voxel size. These bin values represent the diameter of the sphere associated with each marrow-space voxel in the local-thickness calculation, with each bin represented by its mid-range value for histogram-based analysis. The corresponding Tb.Sp maps are exported as 8-bit greyscale images, in which grey level 0 represents the background and non-analysed voxels, while grey levels 1–255 encode the corresponding Tb.Sp histogram bins. The grey level, therefore, functions as a bin index rather than a conventional image-intensity value: grey level 1 represents the smallest local-separation class, corresponding to sphere diameters of 1 – <3 voxels; grey level 2 represents sphere diameters of 3 – <5 voxels; and subsequent grey levels encode progressively larger local-separation classes, while preserving their spatial location within the VOI. For visualisation, these greyscale maps were displayed using a custom-made rainbow transfer function, meaning that each grey-level/bin-index value was assigned a corresponding colour according to a predefined colour scale. This colour coding did not alter the underlying Tb.Sp values, but provided a clearer visualisation of the spatial distribution of low- and high-diameter marrow-space regions. Thus, the Tb.Sp map provides the spatial counterpart to the histogram, allowing the modal structure of the Tb.Sp distribution to be related back to anatomical location within the VOI. These Tb.Sp histograms and maps formed the basis of the subsequent distributional analysis and decomposition.

Maximum trabecular separation, max(Tb.Sp), was also extracted from the CTAn Tb.Sp histogram as an additional descriptor of the upper end of the marrow-space separation distribution. For each sample, max(Tb.Sp) was defined as the largest Tb.Sp bin mid-point with a non-zero marrow-space volume. This metric therefore represents the largest detected local-separation class within the analysed VOI, rather than a volume-weighted average.

### 2.3 Decomposition of trabecular separation into local trabecular separation (Tb.Sp_L_) and marrow cavity separation (Tb.Sp_M_)

To characterise heterogeneity within the marrow-space phase, standard Tb.Sp analysis was extended by separating lower- and higher-diameter components of the voxel-wise Tb.Sp distribution using the workflow summarised in Figure 2. Given that the Tb.Sp maps were exported as 8-bit greyscale images, in which grey levels encode Tb.Sp histogram bins corresponding to sphere-diameter ranges, decomposition was performed by applying a grey-level threshold to the Tb.Sp map dataset.

**Figure 2.**
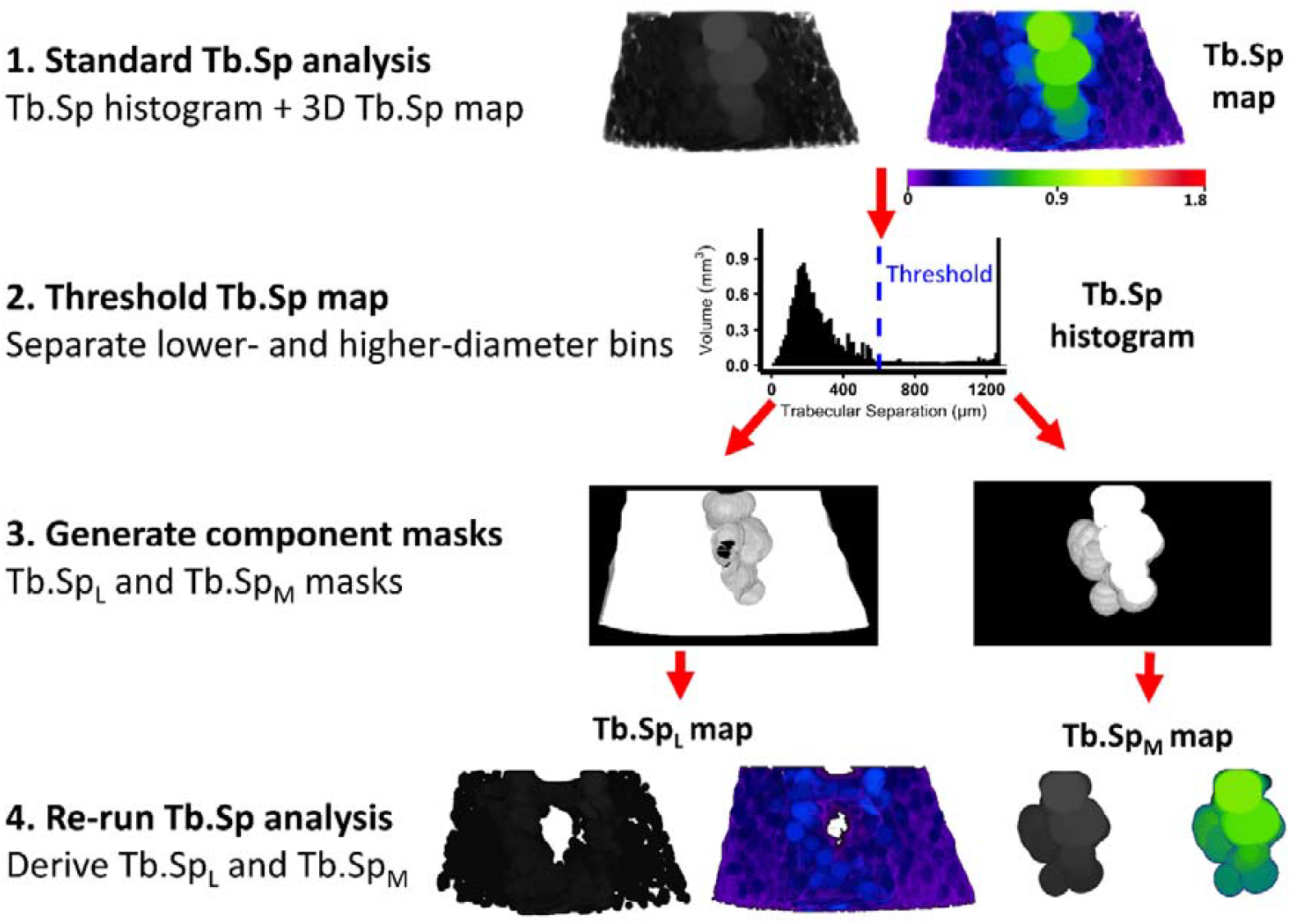
Workflow for decomposition of trabecular separation (Tb.Sp) into local trabecular separation (Tb.Sp_L_) and marrow cavity separation (Tb.Sp_M_). Standard Tb.Sp analysis was first performed in CTAn to generate a Tb.Sp histogram and corresponding 3D voxel-wise Tb.Sp map. A grey-level threshold, corresponding to a selected bin in the Tb.Sp histogram, was then applied to the 3D Tb.Sp map to separate lower- and higher-diameter local-separation values. The thresholded component-specific greyscale Tb.Sp datasets were then binarised to generate lower- and higher-diameter masks, corresponding to the Tb.Sp_L_ and Tb.Sp_M_ regions, respectively. These masks were then applied as component-specific VOIs to the original trabecular dataset, after which standard Tb.Sp analysis was repeated within each VOI to derive Tb.Sp_L_ and Tb.Sp_M_ values. Colour-coded renderings were generated using a rainbow transfer function to visualise the spatial distribution of separation values; this visualisation did not alter the underlying Tb.Sp data.

For each study cohort (OVX, SCI and AGE), metaphyseal Tb.Sp histograms from all specimens, including both control and osteoporotic groups, were reviewed together to assess distribution shape and identify a single separation threshold value corresponding to the boundary between the lower- and higher-diameter components. This threshold was then applied uniformly to all Tb.Sp maps within that cohort. Voxels below the threshold were retained as the lower-diameter Tb.Sp component, whereas voxels at or above the threshold were retained as the higher-diameter Tb.Sp component. This produced two component-specific greyscale Tb.Sp datasets for each specimen. These component-specific greyscale datasets were then binarised to generate two component masks: a lower-diameter mask corresponding to the local trabecular separation region (Tb.Sp_L_), and a higher-diameter mask corresponding to the marrow cavity separation region (Tb.Sp_M_). Because grey-level bin indices correspond to calibrated separation ranges according to voxel size, thresholds are reported both as grey-level values and as approximate calibrated separation values. The higher-diameter Tb.Sp_M_ region was defined using grey-level thresholds of ≥ 50 for OVX and AGE, and ≥ 43 for SCI, corresponding to calibrated separation values of approximately 450 µm, 450 µm, and 600 µm, respectively.

The resulting component masks were used to define subregions within the original metaphyseal VOI, and standard Tb.Sp analysis was repeated within each subregion to obtain Tb.Sp_L_ and Tb.Sp_M_, respectively. Thus, the decomposition was implemented as a mask-based extension of the standard Tb.Sp workflow. This approach preserves the conventional morphometric framework while enabling the lower- and higher-diameter contributions to mean Tb.Sp to be quantified separately.

### 2.4 Distribution analysis and visualisation

To further characterise trabecular structure beyond mean morphometric values, Tb.Th and Tb.Sp histogram outputs from CTAn were analysed quantitatively. Distribution descriptors, including standard deviation and skewness, were calculated from the exported binned histogram data using the bin mid-range values and corresponding bin volumes. These descriptors were used to assess the spread, asymmetry, and shape of the Tb.Th and Tb.Sp distributions. Histogram processing and figure generation were performed using custom Python scripts. For each histogram, bin mid-range values were treated as the representative thickness or separation value for that bin, and bin volumes were used as weights for calculating distribution-level statistics. The Python code is available at [GitHub repository/link to be added].

Voxel-wise Tb.Sp maps generated by CTAn were visualised in CTVox software (version 3.3.1, Bruker, Belgium) using the custom rainbow transfer function described above. These renderings illustrated the spatial distribution of lower- and higher-diameter marrow-space regions and supported interpretation of Tb.Sp_L_ and Tb.Sp_M_.

### 2.5 Comparative analysis of epiphyseal trabecular bone

To assess whether bimodal Tb.Sp distributions were specific to metaphyseal trabecular separation, a comparative analysis was performed using the distal femur SCI dataset, including the representative sample from both SHAM-control and SCI groups. Epiphyseal trabecular bone was manually delineated in CTAn. Standard trabecular morphometric analysis was then performed as described above, and Tb.Sp histograms and voxel-wise maps were exported for comparison with the corresponding metaphyseal Tb.Sp distributions. Tb.Th histograms and maps were also exported from both epiphyseal and metaphyseal VOIs to determine whether bimodality was specific to Tb.Sp or was also present in bone-phase local-thickness distributions.

### 2.6 Statistical analysis

Data are presented as mean ± standard deviation. Statistical analyses were performed in Python using standard scientific computing libraries. A *p*-value of < 0.05 was considered statistically significant.

For group comparisons, conventional trabecular morphometric parameters and Tb.Sp-derived measurements were analysed using the same statistical approach. Conventional parameters included BV/TV, Tb.Th, and Tb.N. Tb.Sp-derived measurements included mean Tb.Sp, Tb.Sp_L_, and Tb.Sp_M_, together with the standard deviation and skewness of each corresponding distribution. For the SCI and OVX datasets, comparisons were performed using unpaired Welch’s t-tests. For the AGE dataset, variables were analysed using one-way ANOVA followed by post hoc pairwise comparisons between groups, with correction for multiple comparisons within each variable. Significance annotations are reported as NS, not significant; * *p* < 0.05; ** *p* < 0.01; *** *p* < 0.001; and **** *p* < 0.0001.

## 3. Results

### 3.1 Trabecular separation distributions are non-Gaussian and multimodal in metaphyseal bone

Metaphyseal Tb.Sp distributions were first examined to determine whether voxel-wise separation values were adequately represented by a single mean. Representative Tb.Sp histograms from metaphyseal trabecular bone VOIs across the SCI, OVX, and AGE cohorts are shown in Figure 3.

**Figure 3.**
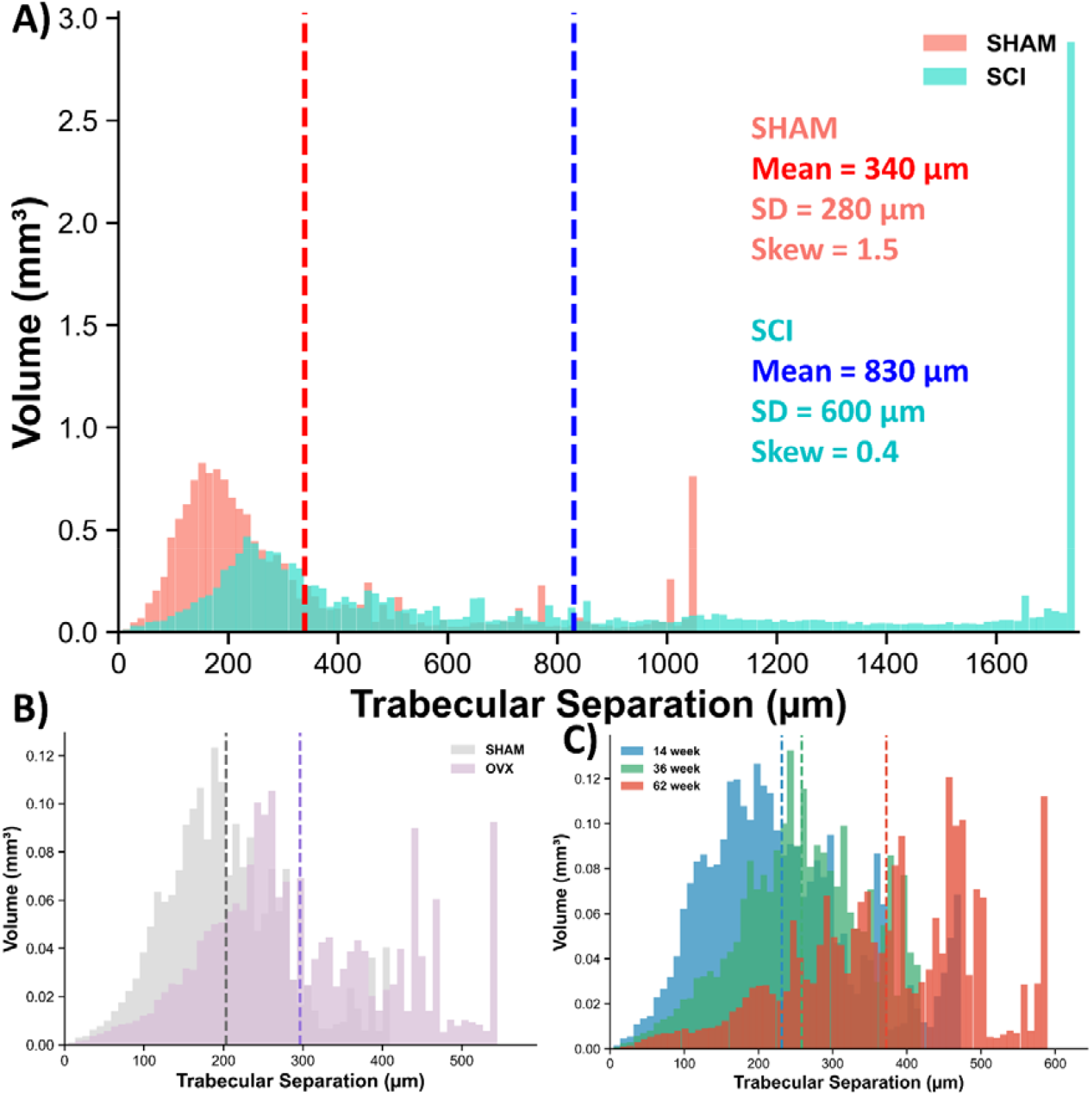
Representative Tb.Sp histograms from metaphyseal trabecular bone VOIs across the SCI, OVX, and AGE cohorts. A) SCI, spinal cord injury; B) OVX, ovariectomy; and C) AGE, ageing. Histograms show raw marrow-space volume per Tb.Sp bin (mm^3^), with dashed vertical lines indicating mean Tb.Sp. Mean, standard deviation, and skewness are additionally shown for the SCI comparison. Across datasets, Tb.Sp distributions were broad and frequently bimodal or multimodal rather than narrow, unimodal, and Gaussian-like.

Across datasets, metaphyseal Tb.Sp distributions were frequently broad, right-skewed, and multimodal rather than approximating a unimodal Gaussian form. This was most pronounced in the SCI dataset, where SCI samples showed a marked shift toward higher separation values and the emergence of a high-diameter peak compared with SHAM controls (Figure 3A).

Similar changes were observed in the OVX and AGE datasets. OVX samples showed broadening of the Tb.Sp distribution relative to SHAM controls (Figure 3B), while AGE samples showed progressive changes in distribution shape with increasing age, including a greater contribution from higher separation values in older animals (Figure 3C). Together, these findings indicate that metaphyseal Tb.Sp distributions contain structural information that is not captured by the conventional mean Tb.Sp alone.

### 3.2 Bimodality is specific to metaphyseal trabecular separation

To determine whether this distributional behaviour was specific to metaphyseal Tb.Sp, we compared metaphyseal Tb.Sp with related local-thickness outputs from the same representative SHAM-control and SCI datasets in Figure 3A. Epiphyseal Tb.Sp was examined to test whether similar multimodality occurred in another trabecular compartment, while epiphyseal and metaphyseal Tb.Th were examined to determine whether multimodality was a general feature of the local-thickness calculation itself (Figure 4).

**Figure 4.**
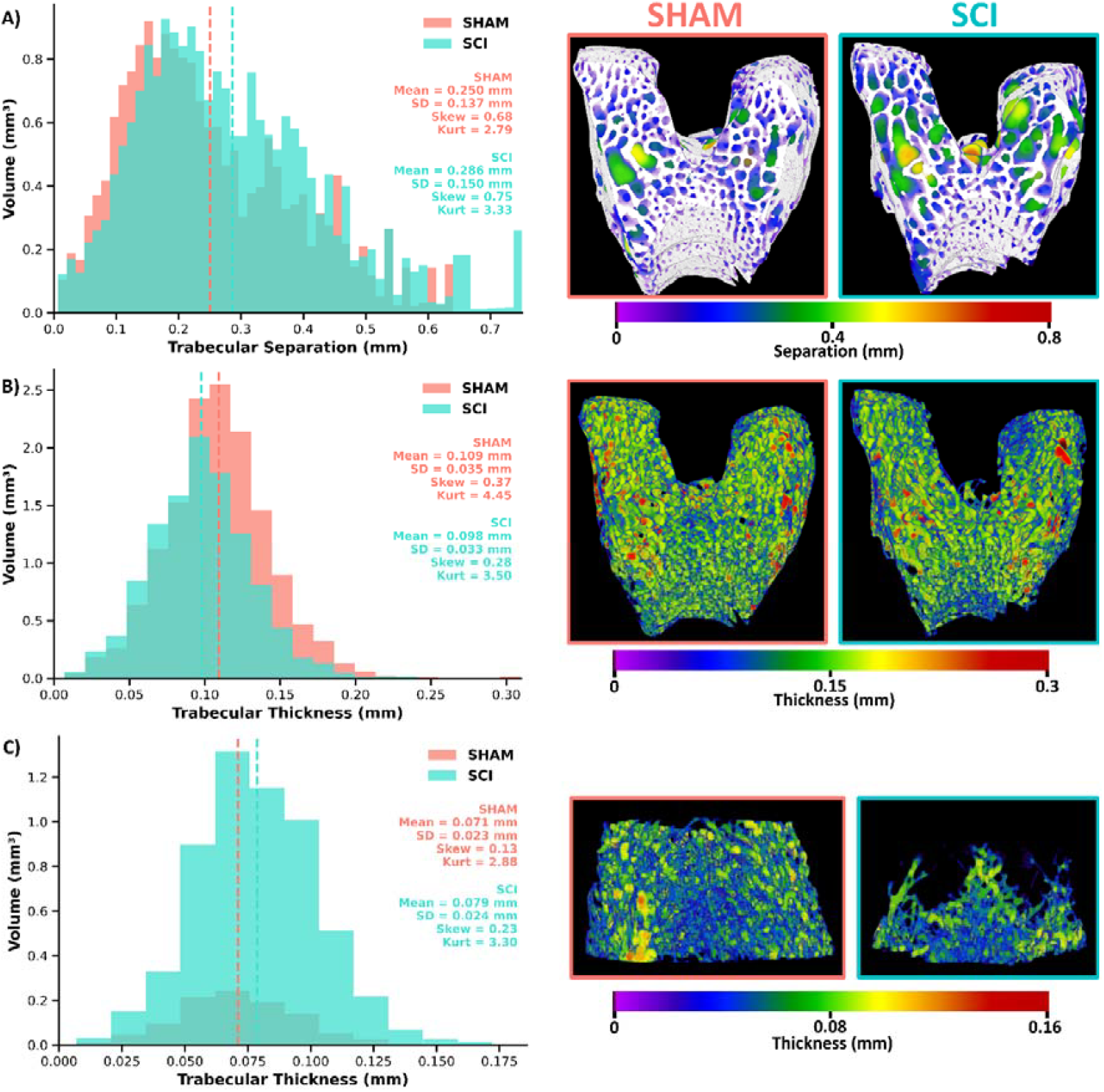
Epiphyseal Tb.Sp and Tb.Th distributions lack the pronounced high-diameter component observed in metaphyseal Tb.Sp. (A) Representative epiphyseal Tb.Sp histograms and maps from SHAM-control and SCI datasets, with binarised trabecular bone shown in white. (B) Representative epiphyseal trabecular thickness (Tb.Th) histograms and maps from the same specimens. (C) Representative metaphyseal Tb.Th histograms and maps. Histograms are shown as raw volume per bin, with dashed vertical lines indicating histogram-weighted mean values. Histogram-derived summary statistics, including mean, standard deviation, skewness, and kurtosis, are shown. Colour scales indicate the local sphere diameter assigned to each voxel by the local-thickness calculation. A custom rainbow transfer function was applied to the Tb.Sp and Tb.Th maps for visualisation.

In contrast to metaphyseal Tb.Sp, epiphyseal Tb.Sp distributions showed a more clearly unimodal profile and lacked the pronounced high-diameter component observed in the metaphyseal region (Figure 4A; compare with Figure 3A). Skewness values did not uniformly distinguish metaphyseal from epiphyseal Tb.Sp, indicating that skewness alone is insufficient to define this behaviour. However, the epiphyseal distributions did not show the same clear separation into lower- and higher-diameter components. The corresponding Tb.Sp maps supported this interpretation, with local separation values distributed across the epiphyseal trabecular space rather than concentrated within a visually dominant high-diameter marrow region.

Trabecular thickness distributions showed a similar contrast with metaphyseal Tb.Sp. Both epiphyseal and metaphyseal Tb.Th histograms were approximately unimodal and lacked the pronounced high-value component observed in metaphyseal Tb.Sp (Figures 4B and 4C). In the corresponding Tb.Th maps, local thickness values were distributed across the trabecular structure without forming a distinct high-value spatial region.

Together, these comparisons indicate that the bimodal/multimodal behaviour observed in metaphyseal Tb.Sp is not simply a universal consequence of voxel-wise local-thickness morphometry. Instead, it appears most evident when the local-thickness approach is applied to the marrow-space phase within metaphyseal trabecular VOIs, consistent with the distinct geometry of the metaphyseal marrow space.

### 3.3 Decomposition of Tb.Sp reveals two distinct structural compartments

Having established that metaphyseal Tb.Sp distributions are frequently non-Gaussian and multimodal, we next examined how these distributional features correspond to spatial organisation within the metaphyseal marrow space. The representative SCI dataset from Figure 3 was selected as a worked example because it exhibited a clearly bimodal Tb.Sp distribution.

The colour-coded metaphyseal Tb.Sp map showed that local separation values were not evenly distributed throughout the VOI (Figure 5B). Lower Tb.Sp values were associated with regions of the remaining trabecular network, whereas higher Tb.Sp values were concentrated within larger contiguous regions of the marrow space. This spatial pattern corresponded closely to the histogram, which contained a lower-diameter component and a higher-diameter component (Figure 5A).

**Figure 5.**
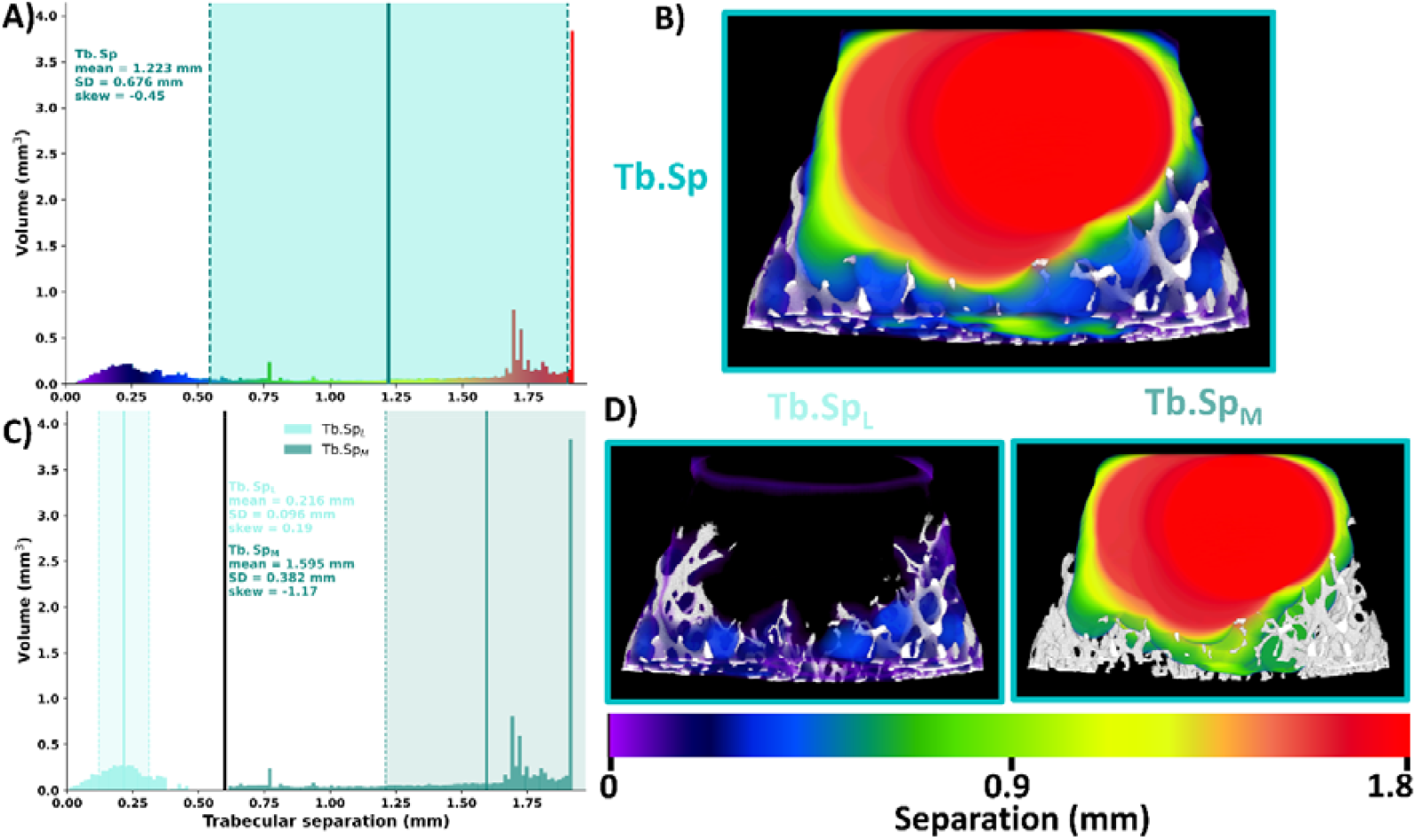
Decomposition of trabecular separation (Tb.Sp) in a representative SCI metaphyseal trabecular bone dataset. **A)** Raw-volume voxel-wise Tb.Sp histogram, with histogram-derived mean, standard deviation (SD), and skewness shown. The solid vertical line indicates mean Tb.Sp, dashed vertical lines indicate ±1 SD, and the transparent shaded region represents the ±1 SD range. **B)** Corresponding voxel-wise Tb.Sp map, with local separation values displayed using a custom rainbow transfer function. **C)** Decomposition of the Tb.Sp distribution into lower- and higher-diameter components, termed local trabecular separation (Tb.Sp_L_) and marrow cavity separation (Tb.Sp_M_), respectively. Histogram-derived mean, SD, and skewness are shown for each component. The black vertical line indicates the threshold used (0.6 mm) to separate Tb.Sp_L_ and Tb.Sp_M_. **D)** Spatial localisation of Tb.Sp_L_ and Tb.Sp_M_ within the same metaphyseal marrow-space map, shown with the same custom rainbow transfer function. Tb.Sp_L_ corresponds to lower-diameter marrow spaces within the trabecular network, whereas Tb.Sp_M_ identifies larger high-diameter marrow-space regions.

A fixed global threshold of 0.6 mm, positioned at the trough between the two histogram components, was then used to decompose the Tb.Sp distribution into local trabecular separation (Tb.Sp_L_) and marrow cavity separation (Tb.Sp_M_) (Figure 5C). Tb.Sp_L_ represented the lower-diameter component of the distribution, while Tb.Sp_M_ represented the higher-diameter component.

When mapped back into the metaphyseal volume, the two components showed distinct spatial localisation (Figure 5D). Tb.Sp_L_ was distributed throughout the trabecular network and corresponded to local spacing between adjacent trabeculae, whereas Tb.Sp_M_ was localised to larger contiguous marrow spaces where trabecular elements were sparse or absent. Recalculating Tb.Sp within these component-specific regions therefore yielded two quantitative measures that retained both the distributional and spatial information contained in the original voxel-wise Tb.Sp map. This provided the basis for comparing Tb.Sp_L_ and Tb.Sp_M_ across osteoporotic models.

### 3.4 Tb.Sp_L_ and Tb.Sp_M_ distinguish patterns of bone loss across models

We next applied the Tb.Sp decomposition approach across the SCI, OVX, and AGE datasets. For each model, standard trabecular morphometric parameters were compared with the mean Tb.Sp, Tb.Sp_L_, and Tb.Sp_M_, and their distributional descriptors. Representative voxel-wise maps were used to relate these quantitative differences to the spatial organisation of the metaphyseal marrow space.

#### 3.4.1 SCI model

Standard trabecular bone morphometry confirmed marked trabecular bone loss in the SCI model. Compared with SHAM-control, SCI animals had lower BV/TV, Tb.Th, and Tb.N, and higher Tb.Sp (Figure 6A–D), consistent with trabecular bone loss and thinning following complete SCI (Williams et al., 2022).

**Figure 6.**
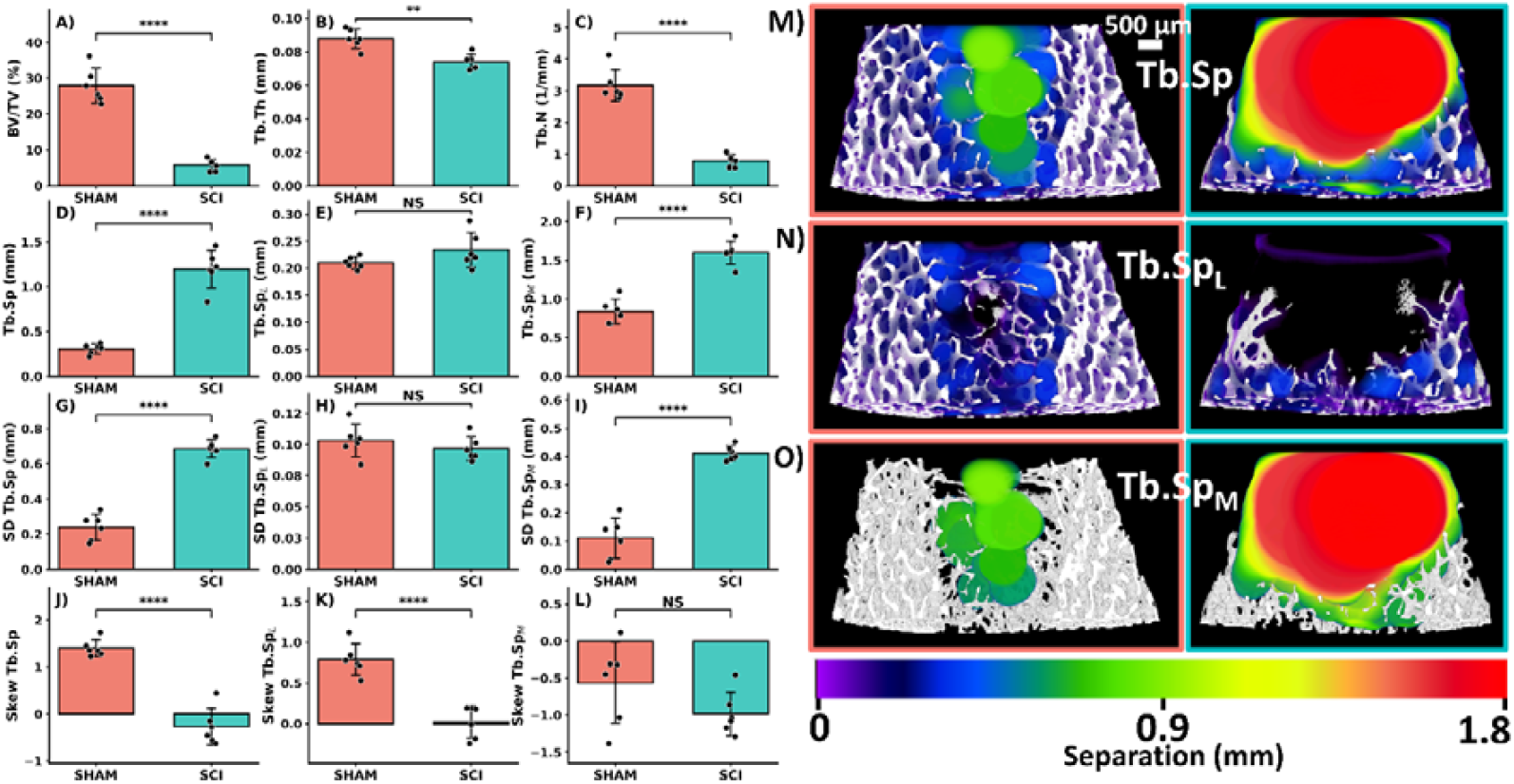
Standard trabecular morphometry and Tb.Sp decomposition at the distal femoral metaphysis in the spinal cord injury (SCI) model. (A–C) Standard trabecular bone morphometric parameters showing BV/TV, Tb.Th and Tb.N in SHAM-control and SCI. (D–F) Tb.Sp and its decomposed lower- and higher-diameter components, termed local trabecular separation (Tb.Sp_L_) and marrow cavity separation (Tb.Sp_M_), respectively. (G–I) Standard deviation (SD) and (J–L) skewness of Tb.Sp, Tb.Sp_L_, and Tb.Sp_M_ distributions. Error bars show mean ± standard deviation with individual data points overlaid. Statistical significance is indicated as shown; NS, not significant. (M-O) Representative voxel-wise maps show the spatial distribution of Tb.Sp, Tb.Sp_L_, and Tb.Sp_M_ in SHAM-control and SCI metaphyseal trabecular bone, displayed using a custom rainbow transfer function where colour represents local sphere diameter/separation magnitude. Scale bar = 500 µm.

Tb.Sp decomposition showed that the increase in mean Tb.Sp was primarily associated with the higher-diameter component, Tb.Sp_M_, whereas Tb.Sp_L_ did not differ significantly between groups (Figure 6D–F). This indicates that the higher mean Tb.Sp in SCI was not driven by a uniform widening of local inter-trabecular spacing, but by increased contribution from larger marrow-space regions.

Distributional metrics supported this interpretation. SCI animals showed a broader overall Tb.Sp distribution, reflected by increased SD(Tb.Sp), with the largest difference observed in SD(Tb.Sp_M_) (Figure 6G–I). Skewness also differed between SHAM-control and SCI for Tb.Sp and Tb.Sp_L_, indicating altered distributional shape (Figure 6J–L).

Representative voxel-wise maps confirmed the spatial basis of these differences (Figure 6M–O). In SCI, high Tb.Sp and Tb.Sp_M_ values were concentrated within large contiguous marrow spaces in the metaphysis, whereas Tb.Sp_L_ remained localised to the residual trabecular network. Together, these findings indicate that SCI-induced trabecular deterioration was dominated by marrow-space expansion, with comparatively limited change in the mean local spacing of the remaining trabecular network.

#### 3.4.2 OVX model

Standard morphometric analysis confirmed trabecular bone loss in the OVX model. Compared with SHAM-control, OVX animals had lower BV/TV and Tb.N, and higher Tb.Sp, while Tb.Th was not significantly different (Figure 7A–D).

**Figure 7.**
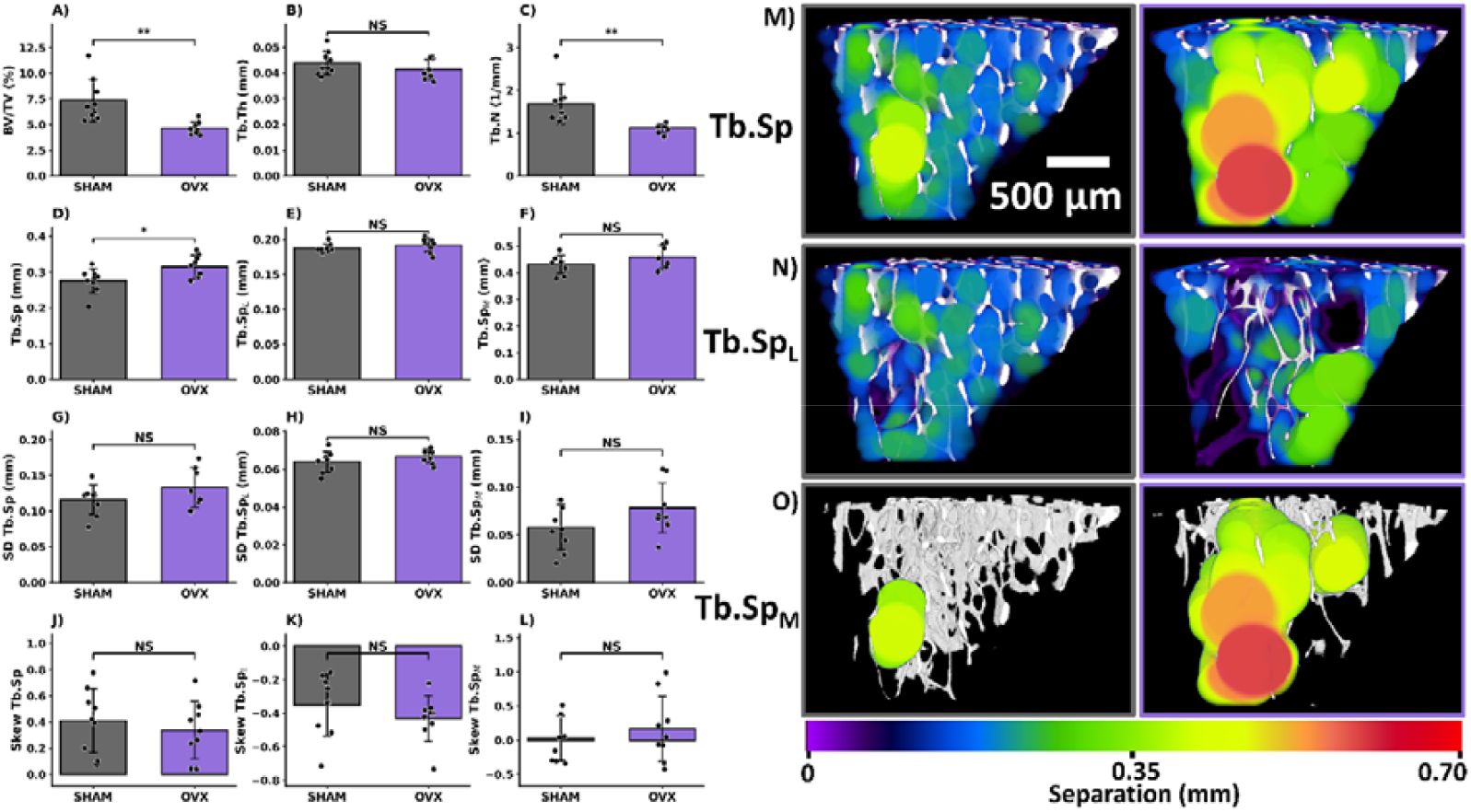
Standard trabecular morphometry and Tb.Sp decomposition at the proximal tibial metaphysis in the ovariectomy (OVX) model. (A–C) Standard trabecular bone morphometric parameters showing BV/TV, Tb.Th and Tb.N in SHAM-control and OVX. (D–F) Tb.Sp and its decomposed lower- and higher-diameter components, termed local trabecular separation (Tb.Sp_L_) and marrow cavity separation (Tb.Sp_M_), respectively. (G–I) Standard deviation (SD) and (J–L) skewness of Tb.Sp, Tb.Sp_L_, and Tb.Sp_M_ distributions. Error bars show mean ± standard deviation with individual data points overlaid. Statistical significance is indicated as shown; NS, not significant. (M-O) Representative voxel-wise maps show the spatial distribution of Tb.Sp, Tb.Sp_L_, and Tb.Sp_M_ in SHAM-control and OVX metaphyseal trabecular bone, displayed using a custom rainbow transfer function where colour represents local sphere diameter/separation magnitude. Scale bar = 500 µm.

In contrast to the SCI model, decomposition did not identify a single dominant component driving the difference in mean Tb.Sp. Neither Tb.Sp_L_ nor Tb.Sp_M_ differed significantly between SHAM-control and OVX animals (Figure 7E,F). This suggests that, in this dataset, the OVX-associated increase in mean Tb.Sp reflected a more distributed alteration of the separation profile rather than a clear expansion of either the local trabecular or marrow-cavity component alone.

The distributional descriptors were also comparatively preserved. SD(Tb.Sp), SD(Tb.Sp_L_), and SD(Tb.Sp_M_) did not differ significantly between groups (Figure 7G–I), and skewness values were similarly unchanged (Figure 7J–L). Thus, although OVX increased the mean Tb.Sp, this was not accompanied by significant changes in distribution width or asymmetry.

Representative maps supported this interpretation (Figure 7M–O). Compared with SCI, the OVX maps showed less evidence of a dominant expansion of the higher-diameter Tb.Sp_M_ compartment, consistent with a subtler spatially distributed change in metaphyseal trabecular architecture.

#### 3.4.3 AGE model

Standard morphometric analysis showed progressive trabecular bone loss with age. BV/TV and Tb.N were lower in the 36- and 62-week groups compared with 14-week animals, whereas Tb.Th did not differ significantly between groups (Figure 8A–C). Mean Tb.Sp increased with age, with the highest values observed in the 62-week group (Figure 8D).

**Figure 8.**
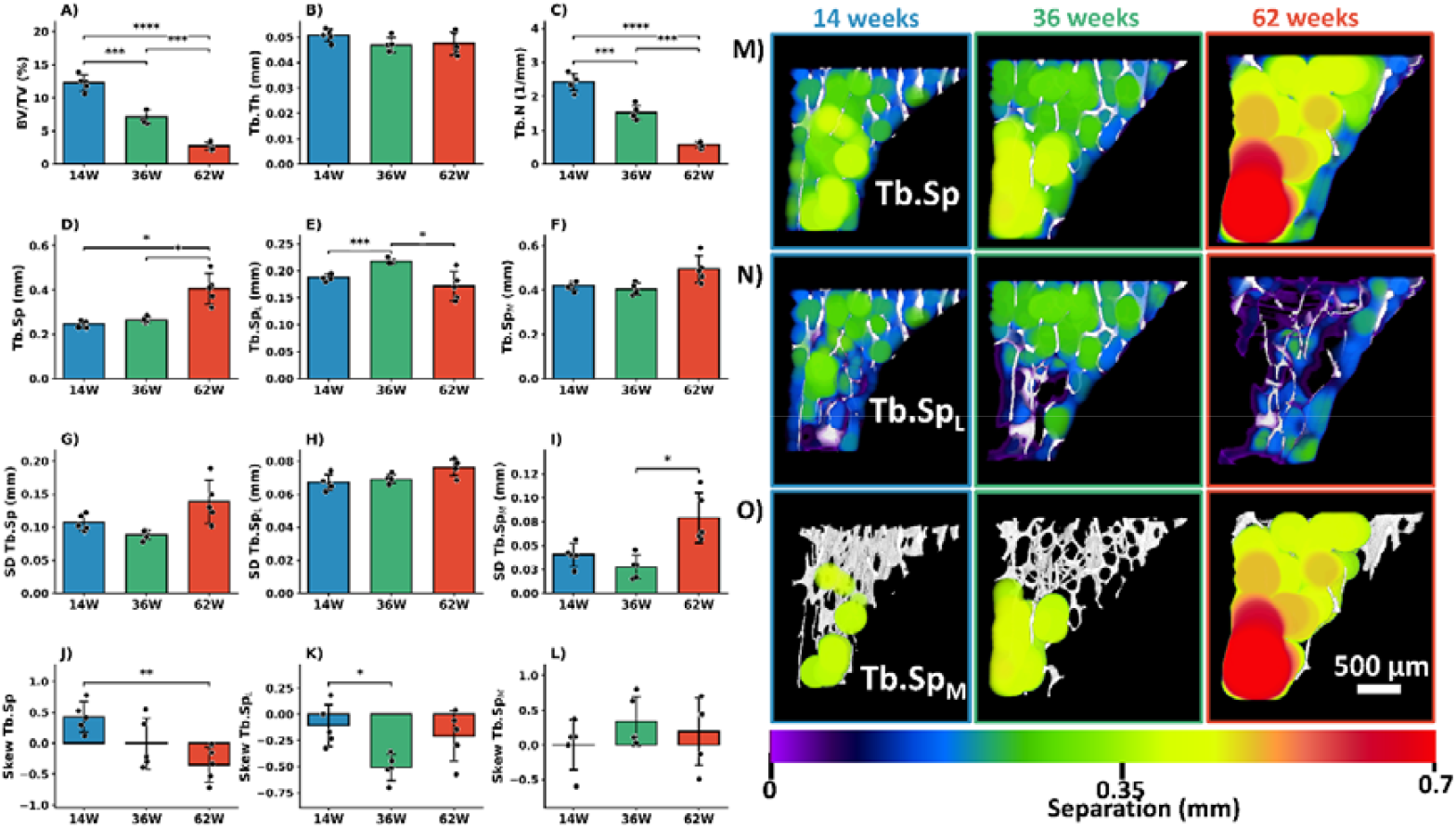
Standard trabecular morphometry and Tb.Sp decomposition at the proximal tibial metaphysis in the ageing model. (A–C) Standard trabecular bone morphometric parameters showing BV/TV, Tb.Th and Tb.N in 14-, 36-, and 62-week-old mice. (D–F) Mean Tb.Sp and its decomposed lower- and higher-diameter components, termed local trabecular separation (Tb.Sp_L_) and marrow cavity separation (Tb.Sp_M_), respectively. (G–I) Standard deviation (SD) and (J–L) skewness of Tb.Sp, Tb.Sp_L_, and Tb.Sp_M_ distributions. Error bars show mean ± standard deviation with individual data points overlaid. Statistical significance is indicated as shown; NS, not significant. (M–O) Representative voxel-wise maps show the spatial distribution of Tb.Sp, Tb.Sp_L_, and Tb.Sp_M_ in 14-, 36-, and 62-week metaphyseal trabecular bone, displayed using a custom rainbow transfer function where colour represents local sphere diameter/separation magnitude. Scale bar = 500 µm.

Tb.Sp decomposition indicated that age-related changes involved both decomposed components, but with different patterns. Tb.Sp_L_ was higher at 36 weeks than at 14 weeks, whereas Tb.Sp_M_ showed its greatest increase in the 62-week group (Figure 8E,F). This suggests that ageing was associated with both altered local spacing within the trabecular network and increasing contribution from larger marrow-space regions in older animals.

Distributional metrics showed corresponding changes in the structure of the Tb.Sp distributions. SD(Tb.Sp_M_) was higher in the 62-week group, indicating greater variability in the higher-diameter marrow-space component (Figure 8G–I). Skewness also differed for Tb.Sp and Tb.Sp_L_, consistent with age-associated changes in distributional shape rather than a simple increase in mean separation alone (Figure 8J–L).

Representative maps illustrated these changes spatially (Figure 8M–O). In 14-week animals, high-diameter Tb.Sp_M_ regions were relatively limited, whereas 62-week animals showed larger and more prominent marrow-space regions. Tb.Sp_L_ remained associated with the trabecular network across age groups. These findings suggest that age-related trabecular deterioration represents a mixed structural phenotype, involving both local changes within the trabecular network and progressive expansion of larger marrow-space regions.

As a supplementary descriptor of the upper tail of the Tb.Sp distribution, max(Tb.Sp) was also extracted from the original Tb.Sp histograms (Supplementary Figure 1). max(Tb.Sp) was higher in SCI than in SHAM-control, consistent with the marked expansion of large marrow-space regions observed in the Tb.Sp_M_ maps. In the OVX model, max(Tb.Sp) did not differ significantly between groups, in agreement with the absence of a significant Tb.Sp_M_ difference. In the AGE dataset, max(Tb.Sp) was highest in the 62-week group, further supporting the emergence of larger marrow-space separations with ageing. These findings were consistent with the main decomposition analysis and support the interpretation that the upper end of the Tb.Sp distribution differs across models of trabecular bone loss.

Taken together, Tb.Sp decomposition revealed different patterns of trabecular separation across the three models. In SCI, increased Tb.Sp was mainly associated with the higher-diameter Tb.Sp_M_ component. In OVX, mean Tb.Sp increased, with no significant differences in Tb.Sp_L_, Tb.Sp_M_, distribution width, or skewness. In ageing, both Tb.Sp_L_ and Tb.Sp_M_ were altered, with the largest increases in marrow-space diameter observed in the oldest animals.

## 4. Discussion

### 4.1 Principal findings

This study demonstrates that trabecular separation (Tb.Sp) in metaphyseal trabecular bone is frequently non-Gaussian and multimodal. Across spinal cord injury (SCI), ovariectomy (OVX), and ageing datasets, Tb.Sp histograms were commonly skewed and contained more than one apparent component, indicating that the underlying marrow-space structure is not adequately represented by a single average value, Tb.Sp, alone.

By decomposing Tb.Sp into lower- and higher-diameter components, termed local trabecular separation (Tb.Sp_L_) and marrow cavity separation (Tb.Sp_M_), we identified two spatially distinct contributions to the overall Tb.Sp measurement. Tb.Sp_L_ was associated mainly with local separation within the residual trabecular network, whereas Tb.Sp_M_ corresponded to larger contiguous marrow-space regions. These components, therefore, provide a distribution-derived description of metaphyseal marrow space, separating local trabecular spacing from larger regions of trabecular loss or cavity expansion.

The relative behaviour of Tb.Sp_L_ and Tb.Sp_M_ differed between models, suggesting that increases in mean Tb.Sp can reflect different structural processes. In SCI, the dominant change was in Tb.Sp_M_, consistent with expansion of larger marrow cavities. In OVX, the absence of significant component-specific or distributional changes despite increased mean Tb.Sp suggests a subtler or more distributed alteration of trabecular spacing. In ageing, changes in both components indicate a mixed phenotype, with the higher-diameter marrow-space component becoming most prominent in older animals.

### 4.2 Implications for trabecular morphometry

Standard trabecular morphometry is commonly summarised using mean values, including BV/TV, Tb.Th, Tb.N, and Tb.Sp (Bouxsein et al., 2010). These parameters remain essential for confirming trabecular bone loss and describing overall bone architecture. However, the present findings show that the mean Tb.Sp can mask differences in how the marrow space is organised, particularly when local trabecular spacing and larger marrow cavities contribute differently to the same overall value. This issue is especially important when the Tb.Sp distribution is bimodal, because the mean value may lie between the lower- and higher-diameter components rather than representing either component directly. In such cases, mean Tb.Sp should be interpreted as a summary of the overall marrow-space distribution, rather than as a specific structural spacing within the trabecular network.

The proposed decomposition provides a simple extension of the standard morphometric workflow. It uses existing local thickness analysis outputs, the Tb.Sp histogram and voxel-wise Tb.Sp map, to extract additional information on the distribution and spatial localisation of marrow-space diameters. In this context, Tb.Sp_L_ and Tb.Sp_M_ should not be viewed as replacements for conventional Tb.Sp, but as complementary descriptors. Mean Tb.Sp remains the primary standard parameter, while Tb.Sp_L_ and Tb.Sp_M_ help explain whether changes in mean Tb.Sp reflects the distributed widening of local trabecular spacing, expansion of larger marrow cavities, or a combination of both. This may be particularly useful in metaphyseal VOIs, where relatively preserved trabecular networks and expanded marrow-space regions can coexist within the same analysed volume.

The maximum Tb.Sp value, a standard output in BoneJ2, provides an additional simple descriptor of the largest marrow-space separation detected within the VOI (Domander et al., 2021). In the present study, the maximum Tb.Sp broadly followed the decomposition results, showing a marked increase in SCI and an age-associated increase in the oldest animals, while no significant difference was observed in the OVX dataset (Supplementary Figure 1).

### 4.3 Distinct osteoporosis models may exhibit different structural modes of trabecular deterioration

The differing behaviour of Tb.Sp_L_ and Tb.Sp_M_ across models suggests that different forms of osteoporosis may be associated with distinct structural modes of trabecular deterioration. In the SCI dataset, bone loss occurred following motor- and sensory-complete spinal cord injury, producing hindlimb paralysis in which the hindlimbs no longer bear weight and instead lie horizontally on the ground (Williams, et al., 2022b). Although SCI-induced osteoporosis may also involve neural and systemic factors (Jiang, Dai, & Jiang, 2006), this model therefore represents a severe form of disuse-associated bone loss with profound loss of normal mechanical loading. Higher mean Tb.Sp in SCI was mainly associated with the higher-diameter Tb.Sp_M_ component, which localised to larger contiguous marrow cavity regions. One plausible explanation is that severe unloading preferentially affects mechanically under-stimulated regions of the metaphysis. Trabecular extent normally decreases with distance from the growth plate (Williams et al., 2020), and centrally located trabeculae farther from the cortical shell may experience lower local strain than trabeculae closer to load-bearing boundaries. Loss of these trabeculae would expand larger central marrow spaces, consistent with the increased contribution of Tb.Sp_M_. This interpretation aligns with mechanostat-based concepts of bone adaptation (Frost, 2003).

In contrast, the OVX dataset represents bone loss driven primarily by oestrogen deficiency, analogous to post-menopausal osteoporosis (Roberts et al., 2019), because oestrogen deficiency acts systemically on bone remodelling and increases bone turnover, OVX-associated deterioration may be expected to be less spatially localised than in a severe unloading model such as SCI (Weitzmann & Pacifici, 2006). Consistent with this interpretation, OVX did not show the same dominant shift toward Tb.Sp_M_ observed in SCI. Instead, changes in Tb.Sp were more balanced, without significant changes in either decomposed component or in distributional descriptors, suggesting more subtle alteration of trabecular spacing, including thinning or loss of trabecular elements throughout the network, rather than preferential expansion of large central marrow cavities.

The AGE dataset showed a mixed pattern, consistent with the multifactorial nature of age-related bone loss, that includes changes in endocrine status, accumulated remodelling imbalance, and habitual mechanical loading over time. Accordingly, age-associated differences were observed in both Tb.Sp_L_ and Tb.Sp_M_, with the largest marrow-space component becoming most prominent in the oldest animals.

These interpretations should be considered hypothesis-generating. The apparent contrast between SCI and OVX may reflect differences in the dominant biological driver of bone loss, as well as in the rate and timing of structural deterioration. SCI-induced unloading may produce rapid trabecular loss and expansion of larger marrow cavities, whereas OVX-related bone loss may develop more gradually through distributed changes in trabecular spacing and remodelling. It is also possible that earlier or later time points within each model would show different relative contributions of Tb.Sp_L_ and Tb.Sp_M_. Because the component-wise and distributional analyses were exploratory and applied to relatively small pre-existing datasets, the statistical comparisons should be interpreted as supporting the structural interpretation of the distributions rather than as definitive evidence of model-specific mechanisms. Nevertheless, the present findings support the concept that similar changes in mean Tb.Sp may arise from different underlying structural patterns. Decomposition of Tb.Sp into Tb.Sp_L_ and Tb.Sp_M_ therefore provides a potential framework for distinguishing bone loss dominated by distributed widening of the trabecular network from bone loss dominated by expansion of larger marrow cavities. The strength of this interpretation should be considered in light of several limitations.

### 4.4 Limitations and future work

First, the datasets were derived from separate pre-existing studies rather than a single experiment designed to compare osteoporosis models directly. Consequently, the SCI, OVX, and AGE datasets differed in µCT acquisition parameters, experimental design, species, sex, and anatomical site: SCI was analysed in the distal femoral metaphysis of male rats, whereas OVX and AGE were analysed in the proximal tibial metaphysis of female and male mice, respectively. Absolute morphometric values should therefore not be directly compared across models. Instead, the cross-model analysis should be interpreted as evidence that non-Gaussian and multimodal Tb.Sp distributions occur across independent datasets and experimental contexts.

Second, the decomposition of Tb.Sp into Tb.Sp_L_ and Tb.Sp_M_ depends on thresholding the voxel-wise Tb.Sp map. In the present study, a global threshold was applied within each dataset to separate lower- and higher-diameter components. Although this provided a consistent basis for comparing groups, the choice of threshold influences the absolute values of Tb.Sp_L_ and Tb.Sp_M_. Future work should therefore evaluate automated approaches for defining this separation.

A further limitation arises from the maximal sphere-fitting method itself. Large marrow cavities are not represented exclusively by large spheres; instead, smaller spheres are also fitted at cavity margins and in regions where larger spheres cannot be accommodated. As a result, some voxels classified as Tb.Sp_L_ may occur at the edges of larger marrow cavities rather than exclusively within the local trabecular network. Tb.Sp_L_ should therefore be interpreted as an approximate, distribution-derived measure of lower-diameter separation, rather than as a perfectly discrete anatomical compartment. Maximum Tb.Sp and Tb.Sp_M_ also require careful interpretation because they represent the upper extreme(s) of the histogram, which may be affected by isolated artefacts, VOI extent, segmentation, etc. A related methodological consideration is that the Tb.Sp_L_ and Tb.Sp_M_ histograms are not direct subdivisions of the original Tb.Sp histogram. Because the voxel-wise Tb.Sp map was thresholded first and Tb.Sp was then recalculated within the resulting component-specific regions, the recombined Tb.Sp_L_ and Tb.Sp_M_ histograms do not necessarily reproduce the original Tb.Sp histogram exactly. Furthermore, Tb.Sp was calculated in CTAn software, so the exact implementation of the local thickness algorithm is not fully known. However, the proposed decomposition is based on voxel-wise Tb.Sp maps and histograms, and could be readily implemented using open-source tools such as BoneJ2 (Domander et al., 2021). Despite these limitations, the present findings suggest several practical steps for improving the interpretation of Tb.Sp in metaphyseal trabecular bone.

Further future work will extend the analysis beyond sphere-derived local separation values. The maximal sphere-fitting approach underlying Tb.Sp provides a useful and established measure of local marrow-space diameter, but trabecular voids are often anisotropic, irregular, and interconnected. Future work should also examine whether similar distributional behaviour occurs in other trabecular-rich skeletal compartments, including vertebral bodies (Roux et al., 2010), iliac crest biopsies (Chappard et al., 2008), and the calcaneus (Metcalf et al., 2018), as well as clinically important peripheral sites assessed by HR-pQCT, such as the distal radius and distal tibia (Zhu et al., 2016). Alternative geometric descriptions, such as ellipsoid-based methods (Doube, 2015; Felder et al., 2021), can be applied to the marrow space and may provide additional information on the shape and orientation of larger cavities. Similarly, treating the marrow space as a connected pore network could enable quantification of cavity connectivity, pore–throat relationships, transport pathways, and spatial organisation (Dong & Blunt, 2009; Gostick et al., 2016). These approaches may complement Tb.Sp_L_ and Tb.Sp_M_ by describing not only the size of marrow-space compartments, but also their shape and connectivity.

More broadly, this approach may have relevance beyond bone morphometry. µCT is commonly used to quantify pore-size distributions in porous biomaterials and engineered materials, including tissue-engineering scaffolds, ceramic bone substitutes, calcium phosphate cements, polymer foams, 3D-printed lattices, filters, rocks, and soils. These materials often contain mixed pore populations, including small interconnected pores alongside larger voids, channels, defects, or collapsed regions. In such cases, a single mean pore-size value may be difficult to interpret because it may lie between structurally distinct pore populations.

### 4.5 Practical recommendations

Based on these findings, we recommend that future analyses of metaphyseal trabecular bone continue to report standard morphometric parameters, including mean Tb.Sp, but also make greater use of the distributional information already generated during Tb.Sp analysis.

At a minimum, where Tb.Sp histograms are available, studies should consider reporting:

1. mean Tb.Sp, as the standard morphometric parameter;
2. SD(Tb.Sp), to capture distributional width or heterogeneity;
3. maximum Tb.Sp, to describe the largest local marrow-space separation within the VOI;
4. representative Tb.Sp histograms and/or voxel-wise maps where interpretation depends on spatial heterogeneity.

Where histograms are clearly skewed, broad, or multimodal, decomposition into Tb.Sp_L_ and Tb.Sp_M_ may provide additional insight. These values can help distinguish between local widening of the trabecular network and expansion of larger marrow-space cavities. However, the method should be applied with transparent reporting of the thresholding approach, because component values may depend on how the lower- and higher-diameter regions are defined.

Thus, we propose a tiered reporting approach. Standard Tb.Sp should remain the core metric. SD(Tb.Sp) and maximum Tb.Sp provide simple distributional descriptors that are easy to extract from existing histogram outputs. Tb.Sp_L_ and Tb.Sp_M_ provide a more detailed analysis when the biological question concerns spatially heterogeneous bone loss or when the histogram indicates that mean Tb.Sp is unlikely to adequately summarise the distribution.

## Conclusion

Metaphyseal Tb.Sp is frequently non-Gaussian and multimodal, meaning that the mean Tb.Sp may represent an intermediate summary value rather than a specific structural spacing within the trabecular network. Decomposition into Tb.Sp_L_ and Tb.Sp_M_ separates lower-diameter local separation from higher-diameter marrow cavity expansion, revealing spatial and distributional information not captured by mean Tb.Sp alone. This approach provides a simple extension of standard trabecular morphometry and may improve the interpretation of heterogeneous trabecular bone loss, particularly when Tb.Sp histograms are skewed or multimodal.

## CRediT authorship contribution statement

CH: Resources, Writing – review and editing. JCL: Resources, Writing – review and editing. CSG: Resources, Writing – review and editing. JAW: Conceptualisation, Data Curation, Formal Analysis, Investigation, Methodology, Project Administration, Validation, Visualization, Writing - original draft, Writing – review and editing.

## Acknowledgements

No external funding was received for this work.

## References

Bouxsein, M. L., Boyd, S. K., Christiansen, B. A., Guldberg, R. E., Jepsen, K. J., & Müller, R. (2010). Guidelines for assessment of bone microstructure in rodents using micro-computed tomography. Journal of Bone and Mineral Research, 25(7), 1468–1486. 10.1002/jbmr.141

Buie, H. R., Campbell, G. M., Klinck, R. J., MacNeil, J. A., & Boyd, S. K. (2007). Automatic segmentation of cortical and trabecular compartments based on a dual threshold technique for in vivo micro-CT bone analysis. Bone, 41(4), 505–515. 10.1016/j.bone.2007.07.007

Chappard, C., Marchadier, A., & Benhamou, C. L. (2008). Side-to-side and within-side variability of 3D bone microarchitecture by conventional micro-computed tomography of paired iliac crest biopsies. Bone, 43(1), 203–208. 10.1016/j.bone.2008.02.019

Domander, R., Felder, A. A., & Doube, M. (2021). BoneJ2 - refactoring established research software. Wellcome Open Research, 6. 10.12688/wellcomeopenres.16619.1

Dong, H., & Blunt, M. J. (2009). Pore-network extraction from micro-computerized-tomography images. Physical Review E, 80(3), 36307. 10.1103/PhysRevE.80.036307

Doube, M. (2015). The ellipsoid factor for quantification of rods, plates, and intermediate forms in 3D geometries. Frontiers in Endocrinology, 6(FEB). 10.3389/fendo.2015.00015

Erben, R. G. (1996). Trabecular and endocortical bone surfaces in the rat: Modeling or remodeling? The Anatomical Record, 246(1), 39–46. 10.1002/(SICI)1097-0185(199609)246:1<39::AID-AR5>3.0.CO;2-A

Fazzalari, N. L., & Parkinson, I. H. (1996). Fractal dimension and architecture of trabecular bone. The Journal of Pathology, 178(1), 100–105. 10.1002/(SICI)1096-9896(199601)178:1<100::AID-PATH429>3.0.CO;2-K

Felder, A. A., Monzem, S., De Souza, R., Javaheri, B., Mills, D., Boyde, A., & Doube, M. (2021). The plate-to-rod transition in trabecular bone loss is elusive. Royal Society Open Science, 8(6). 10.1098/rsos.201401

Frost, H. M. (2003). Bone’s Mechanostat: A 2003 Update. Anatomical Record - Part A Discoveries in Molecular, Cellular, and Evolutionary Biology, Vol. 275, pp. 1081–1101. Wiley-Liss Inc. 10.1002/ar.a.10119

Gabet, Y., Kohavi, D., Kohler, T., Baras, M., Müller, R., & Bab, I. (2008). Trabecular bone gradient in rat long bone metaphyses: Mathematical modeling and application to morphometric measurements and correction of implant positioning. Journal of Bone and Mineral Research, 23(1), 48–57. 10.1359/jbmr.070901

Gostick, J., Aghighi, M., Hinebaugh, J., Tranter, T., Hoeh, M. A., Day, H., … Putz, A. (2016). OpenPNM: A Pore Network Modeling Package. Computing in Science & Engineering, 18(4), 60–74. 10.1109/MCSE.2016.49

Harrigan, T. P., & Mann, R. W. (1984). Characterization of microstructural anisotropy in orthotropic materials using a second rank tensor. Journal of Materials Science, 19(3), 761–767. 10.1007/BF00540446

Herbst, E. C., Felder, A. A., Evans, L. A. E., Ajami, S., Javaheri, B., & Pitsillides, A. A. (2021). A new straightforward method for semi-automated segmentation of trabecular bone from cortical bone in diverse and challenging morphologies. Royal Society Open Science, 8(8), 210408. 10.1098/rsos.210408

Hildebrand, T., & Rüegsegger, P. (1997). A new method for the model-independent assessment of thickness in three-dimensional images. Journal of Microscopy, 185(1), 67–75. 10.1046/j.1365-2818.1997.1340694.x

Huesa, C., McGrath, S., Dunning, L., Vieri, M. L., McCulloch, K., McIntosh, K. A., … Goodyear, C. S. (2026). PAR2 deletion in the osteoblast lineage affords long-term cartilage protection in experimental osteoarthritis. Osteoarthritis and Cartilage, 34(6), 819–831. 10.1016/j.joca.2026.03.004

Jiang, S.-D., Dai, L.-Y., & Jiang, L.-S. (2006). Osteoporosis after spinal cord injury. Osteoporosis International, 17(2), 180–192. 10.1007/s00198-005-2028-8

Kerckhofs, G., Durand, M., Vangoitsenhoven, R., Marin, C., Van Der Schueren, B., Carmeliet, G., … Vandamme, K. (2016). Changes in bone macro-and microstructure in diabetic obese mice revealed by high resolution microfocus X-ray computed tomography. Scientific Reports, 6. 10.1038/srep35517

Koh, N. Y. Y., Miszkiewicz, J. J., Fac, M. L., Wee, N. K. Y., & Sims, N. A. (2024). Preclinical Rodent Models for Human Bone Disease, Including a Focus on Cortical Bone. Endocrine Reviews, 45(4), 493–520. 10.1210/endrev/bnae004

LeBoff, M. S., Greenspan, S. L., Insogna, K. L., Lewiecki, E. M., Saag, K. G., Singer, A. J., & Siris, E. S. (2022). The clinician’s guide to prevention and treatment of osteoporosis. Osteoporosis International, 33(10), 2049–2102. 10.1007/s00198-021-05900-y

Liu, Z., Yan, C., Kang, C., Zhang, B., & Li, Y. (2015). Distributional variations in trabecular architecture of the mandibular bone: An in vivo micro-CT analysis in rats. PLoS ONE, 10(1). 10.1371/journal.pone.0116194

Metcalf, L. M., Dall’Ara, E., Paggiosi, M. A., Rochester, J. R., Vilayphiou, N., Kemp, G. J., & McCloskey, E. V. (2018). Validation of calcaneus trabecular microstructure measurements by HR-pQCT. Bone, 106, 69–77. 10.1016/j.bone.2017.09.013

Odgaard, A., & Gundersen, H. J. G. (1993). Quantification of connectivity in cancellous bone, with special emphasis on 3-D reconstructions. Bone, 14(2), 173–182. 10.1016/8756-3282(93)90245-6

Porrelli, D., Abrami, M., Pelizzo, P., Formentin, C., Ratti, C., Turco, G., … Murena, L. (2022). Trabecular bone porosity and pore size distribution in osteoporotic patients – A low field nuclear magnetic resonance and microcomputed tomography investigation. Journal of the Mechanical Behavior of Biomedical Materials, 125. 10.1016/j.jmbbm.2021.104933

Reznikov, N., Chase, H., Ben Zvi, Y., Tarle, V., Singer, M., Brumfeld, V., … Weiner, S. (2016). Intertrabecular angle: A parameter of trabecular bone architecture in the human proximal femur that reveals underlying topological motifs. Acta Biomaterialia, 44, 65–72. 10.1016/j.actbio.2016.08.040

Roberts, B. C., Giorgi, M., Oliviero, S., Wang, N., Boudiffa, M., & Dall’Ara, E. (2019). The longitudinal effects of ovariectomy on the morphometric, densitometric and mechanical properties in the murine tibia: A comparison between two mouse strains. Bone, 127, 260–270. 10.1016/j.bone.2019.06.024

Roux, J. P., Wegrzyn, J., Arlot, M. E., Guyen, O., Delmas, P. D., Chapurlat, R., & Bouxsein, M. L. (2010). Contribution of trabecular and cortical components to biomechanical behavior of human vertebrae: An ex vivo study. Journal of Bone and Mineral Research, 25(2), 356–361. 10.1359/jbmr.090803

Salmon, P. (2020). Application of Bone Morphometry and Densitometry by X-Ray Micro-CT to Bone Disease Models and Phenotypes. In K. Orhan (Ed.), Micro-computed Tomography (micro-CT) in Medicine and Engineering (pp. 49–75). Cham: Springer International Publishing. 10.1007/978-3-030-16641-0_5

Salmon, P. L., Monzem, S., Javaheri, B., Oste, L., Kerckhofs, G., & Pitsillides, A. A. (2023). Resolving trabecular metaphyseal bone profiles downstream of the growth plate adds value to bone histomorphometry in mouse models. Frontiers in Endocrinology, 14. 10.3389/fendo.2023.1158099

Seeman, E., & Delmas, P. D. (2006). Bone Quality — The Material and Structural Basis of Bone Strength and Fragility. New England Journal of Medicine, 354(21), 2250–2261. 10.1056/NEJMra053077

Weitzmann, M. N., & Pacifici, R. (2006). Estrogen deficiency and bone loss: an inflammatory tale. The Journal of Clinical Investigation, 116(5), 1186–1194. 10.1172/JCI28550

Williams, J. A., Huesa, C., Turunen, M. J., Oo, J. A., Radzins, O., Gardner, W., … Coupaud, S. (2022). Time course changes to structural, mechanical and material properties of bone in rats after complete spinal cord injury. J Musculoskelet Neuronal Interact, 22(2), 212–234.

Williams, J. A., Huesa, C., Windmill, J. F. C., Purcell, M., Reid, S., Coupaud, S., & Riddell, J. S. (2022). Spatiotemporal responses of trabecular and cortical bone to complete spinal cord injury in skeletally mature rats. Bone Reports, 16(101592). 10.1016/j.bonr.2022.101592

Williams, J. A., Windmill, J. F. C., Tanner, K. E., Riddell, J. S., & Coupaud, S. (2020). Global and site-specific analysis of bone in a rat model of spinal cord injury-induced osteoporosis. Bone Reports, 12(100233). 10.1016/j.bonr.2019.100233

Zhu, T. Y., Hung, V. W. Y., Cheung, W. H., Cheng, J. C. Y., Qin, L., & Leung, K. S. (2016). Value of Measuring Bone Microarchitecture in Fracture Discrimination in Older Women with Recent Hip Fracture: A Case-control Study with HR-pQCT. Scientific Reports, 6. 10.1038/srep34185

