## Supplementary Material for "Beyond mean trabecular separation: decomposing metaphyseal marrow space into local trabecular spacing and marrow cavity expansion"

Department of Biomedical Engineering

University of Strathclyde

Wolfson Building

106 Rottenrow East

GLASGOW

G4 0NW,

United Kingdom

**Contents**

**Supplementary Table 1 - Summary table of experimental datasets, µCT acquisition, and Tb.Sp decomposition parameters.**

**Supplementary Figure 1 - Maximum trabecular separation across experimental datasets**

**Supplementary Table 1.** Summary of biological model, skeletal site, µCT acquisition, image processing, standard trabecular morphometry, and Tb.Sp decomposition parameters for the SCI, OVX, and AGE datasets.

| **Section** | **Parameter** | **SCI dataset** | **OVX dataset** | **AGE dataset** |
| --- | --- | --- | --- | --- |
| **Biological model and experimental design** | **Biological model** | Spinal cord injury | Ovariectomy | Ageing |
|  | **Origin of Data set** | T9 transection of the spinal cord | Contralateral leg of **destabilisation of the medial meniscus** model of osteoarthritis (DMM). | Contralateral leg of Sham (DMM) operated mice |
|  | **Main biological driver** | Paralysis induced mechanical unloading | Oestrogen deficiency | Age-related bone loss |
|  | **Species** | Rat | Mouse | mouse |
|  | **Strain** | Wistar | C57Bl6/J | C57Bl6/J |
|  | **Sex** | Male | Female | Male |
|  | **Age at intervention (weeks)** | 7 | 10.7 | N/A |
|  | **Age at endpoint / scan (weeks)** | 17 | 15.4 | 14, 36 &62 |
|  | **Experimental groups** | SHAM, SCI | SHAM, OVX | 14W, 36W, 62W |
|  | **n per group** | 6 | 9 | 5 |
|  | **Time after intervention** | 10 weeks | 5 weeks after OVX surgery  2 weeks after DMM surgery | N/A |
| **Skeletal site and VOI definition** | **Bone analysed** | Femur | Tibia | Tibia |
|  | **Anatomical site** | Distal femoral metaphysis | Proximal tibial metaphysis | Proximal tibial metaphysis |
|  | **Region analysed** | Metaphyseal trabecular bone | Metaphyseal trabecular bone | Metaphyseal trabecular bone |
|  | **VOI start position** | 0.8 mm from growth plate | 0.4 mm from growth plate | 0.4 mm from growth plate |
|  | **VOI height / depth** | 2.8 mm | 1.5 mm | 2 mm |
| **µCT acquisition and reconstruction** | **µCT scanner** | Bruker Skyscan 1172 | Bruker Skyscan 1272 | Bruker Skyscan 1272 |
|  | **Voxel size (µm)** | 6.9 | 4.5 | 4.5 |
|  | **X-ray voltage (kV)** | 70 | 50 | 50 |
|  | **X-ray current (µA)** | 100 | 200 | 200 |
|  | **Filter** | Al, 0.5 mm | Al, 0.5 mm | Al, 0.5 mm |
|  | **Exposure time (ms)** | 470 | 2250 | 2250 |
|  | **Rotation step (°)** | 0.4 | 0.3 | 0.3 |
|  | **Frame Averaging** | 2 | OFF | OFF |
|  | **Reconstruction software** | NRecon (1.6.9.18) | NRecon (1.7.5.1) | NRecon (1.6.10.2) |
|  | **Beam hardening correction (%)** | 40 | 38 | 38 |
|  | **Ring artefact correction** | 13 | 5 | 5 |
|  | **Smoothing** | 2 | 1 | 1 |
| **Image processing** | **Analysis software** | CTAn v1.23.02 | CTAn v1.23.02 | CTAn v1.23.02 |
|  | **Segmentation method** | Automated (MN008) | Automated (MN008) | Automated (MN008) |
|  | **Threshold range** | 90-255 | 92-255 |  |
|  | **Standard morphometric parameters** | BV/TV, Tb.Th, Tb.N, Tb.Sp | BV/TV, Tb.Th, Tb.N, Tb.Sp | BV/TV, Tb.Th, Tb.N, Tb.Sp |
|  | **Additional morphometric parameters** | max Tb.Sp | max Tb.Sp | max Tb.Sp |
| **Tb.Sp distribution and decomposition analysis** | **Tb.Sp outputs used** | Histogram and voxel-wise map | Histogram and voxel-wise map | Histogram and voxel-wise map |
|  | **Histogram statistics calculated** | Mean, SD, skewness, max Tb.Sp | Mean, SD, skewness, max Tb.Sp | Mean, SD, skewness, max Tb.Sp |
|  | **Tb.Sp map visualisation** | Custom rainbow transfer function | Custom rainbow transfer function | Custom rainbow transfer function |
|  | **Threshold used to define Tb.SpL/Tb.SpM** | 900 µm | 450 µm | 450 µm |
|  | **Statistical tests** | Welch’s t-test | Welch’s t-test | One-way ANOVA with post-hoc comparisons |

**Supplementary Figure 1. Maximum trabecular separation across experimental datasets**

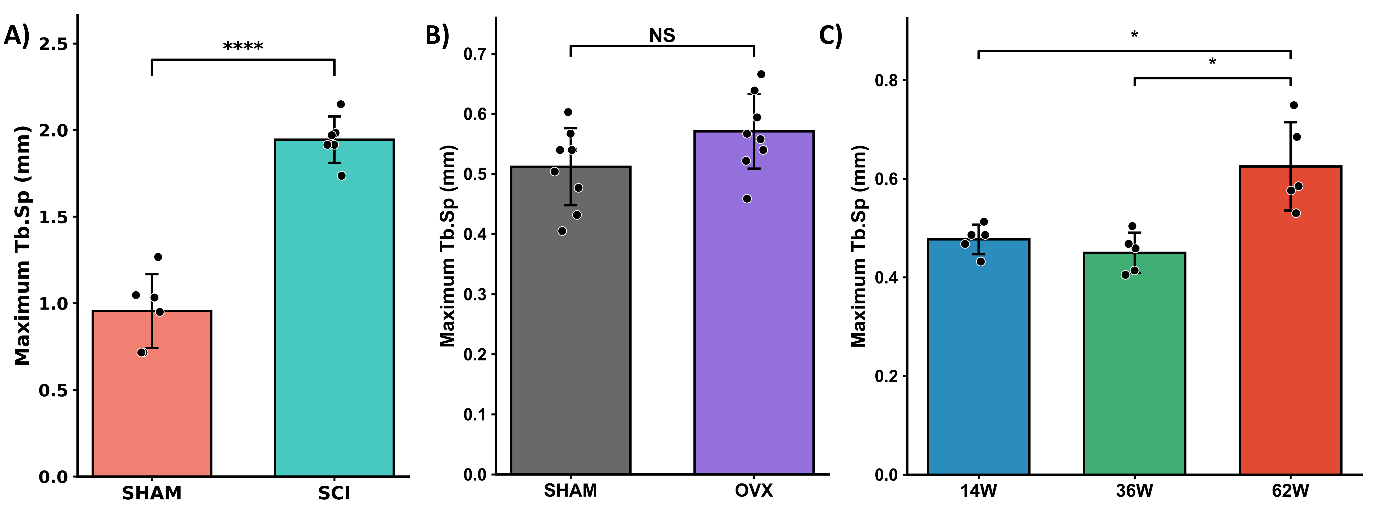

**Supplementary Figure 1.** **Maximum trabecular separation across SCI, OVX, and AGE cohorts.** Maximum trabecular separation (maximum Tb.Sp) was extracted from voxel-wise Tb.Sp histogram exports for each specimen and compared between experimental groups. A) Rat spinal cord injury model: SHAM-control and SCI. B) Mouse ovariectomy model: SHAM-control and OVX. C) Mouse ageing cohort: 14-week, 36-week, and 62-week groups. Bars show mean ± standard deviation with individual specimens overlaid. Statistical significance is indicated as shown; NS, not significant; *p < 0.05; ****p < 0.0001. Maximum Tb.Sp was increased in SCI compared with SHAM-control and in 62-week mice compared with younger age groups, whereas no significant difference was observed between SHAM-control and OVX groups.
